# xiView: A common platform for the downstream analysis of Crosslinking Mass Spectrometry data

**DOI:** 10.1101/561829

**Authors:** Martin Graham, Colin Combe, Lars Kolbowski, Juri Rappsilber

**Author notes:** equal contribution (Martin Graham, Colin Combe).

## Abstract

xiView provides a common platform for the downstream analysis and visualisation of Crosslinking Mass Spectrometry data. It is independent of the search software used and its input is compliant with the relevant mass spectrometry data standards. It uses established visualisation techniques, notably Multiple Coordinated Views, to help the user explore the data and is designed to facilitate comparisons between different datasets.

## Main

Crosslinking Mass Spectrometry (CLMS) investigates the structure and function of proteins by providing proximity information for pairs of amino acids^1^. xiView is a web-based tool for exploring these datasets, as are XLink-DB^2^, xVis^3^, ProXL^4^ and CLMSVault^5^. However, xiView is distinguished from these by (i) standards compliant input data, and (ii) simultaneous interactive multiple views that are linked together, each giving a different yet synchronised perspective on the dataset.

The supported data standards are HUPO-PSI mzIdentML^6^ and HUPO-PSI mzML^7^, with two non-standard formats also supported - a peak list format (.mgf^8^) and one for identifications (CSV). The software has been proven to work with valid mzIdentML, using output from SIM-XL^17^ as an example.

The typical graphical representations of data to be found in both CLMS publications and CLMS software are: a) annotated spectra; b) 2D network representations such as XiNet^9^ and Circos^10^-style views; c) 3D structures with crosslinks mapped onto them; and d) histograms of link distances derived from the 3D structure. xiView expands on this collection with e) a matrix / contact map for chosen protein pairs; f) a protein information panel showing a protein sequence as a wrapped string with crosslinks marked within it; g) visual sequence alignments between search sequences and the chosen PDB file which are partially adjustable; h) a scatterplot; and i) a general-purpose non-distance histogram. Both the scatterplot and histogram display metadata about peptide spectrum matches (PSMs) and/or crosslinks (unique distance restraints) - the specific attributes are selected by the user. All the views along with supporting legends can be exported to figure-quality SVG (Scalable Vector Graphics) files or PNG in the case of the 3D view.

Unlike similar tools, xiView combines all these representations in a single web page and they are all accessible simultaneously. Colour schemes can be chosen based on crosslink attributes and are applied across all views. Crosslinks are filtered using conditions set in a filter bar with the resulting set of crosslinks immediately reflected across all views. Crosslinks selected or highlighted in one view are automatically selected or highlighted in all other open views. This design is in-keeping with the established data visualisation technique known as ‘Multiple Coordinated Views’^11^ - each individual view accentuates and reveals a different aspect of the data, and the coordination between them aids in understanding the relationships between these aspects, which ultimately leads to a better understanding of the dataset.

Across the crosslinking community, it is accepted that *“the term ‘cross-link’ refers to the specific amino acid residues that are connected, irrespective of different peptide sequences due to missed cleavage sites or modifications.”*^12^ A PSM provides a single distance restraint by identifying a pair of crosslinked peptides. Typically, this has been in one spectrum, though isotope labelling and multiple fragmentation techniques can mean a PSM is associated with more than one spectrum (see Section 5.2.9 of the MzIdentML 1.2 specification). There may be several different interpretations of the same spectrum, though search software generally only allows one interpretation of the same spectrum to pass its validation threshold.

xiView uses xiSPEC^13^ for the display of annotated spectra, including the option to view alternative explanations of the same spectrum. Like XLink-DB^2^ (v2) and CLMSVault^5^, xiView uses xiNET^9^ to display a 2D network of the crosslinks. xiNET and the other network views in xiView can also represent ambiguities that result when a peptide occurs in more than one location in the search sequence database.

Uniprot annotations are automatically acquired when search proteins are assigned a valid UniProtKB accession number. A number of types of metadata can also be loaded into xiView. When a PDB^14^ structure is loaded the crosslinks are aligned onto the 3D structures and shown using NGL^15^. The matrix view also shows distances derived from the structure as a shaded background contact map. Often the search sequences do not match the canonical Uniprot sequences, and PDB sequences often offer only partial coverage of sequences. Integrating the Uniprot and PDB data thus requires the calculation of sequence alignments, performed internally within xiView, which are then used to ensure annotations in the xiNet and circular views, and crosslinks and distance measurements in the NGL component, are mapped to the correct place on proteins.

A typical session with xiView is then as follows: an individual or set of crosslink datasets are chosen by a user to view - xiView allows multiple datasets to be aggregated and/or compared through assigning group identifiers. Filtered crosslinks are shown in the collection of co-ordinated views. When crosslinks within one of these views are selected, the corresponding representations of those crosslinks are then accented in the other views to reveal possible patterns e.g. an area of the scatterplot view can be selected to pick crosslinks with a certain number of PSMs and scores for those PSMs, and the selection seen in the circular or network view to see if there is a positional pattern. At the same time, the supporting PSMs and data are shown in a table, and selecting an individual PSM within that table then shows the corresponding annotated spectrum in another view, see Figure 1. Then, using the metadata loading option, an appropriate PDB file can be loaded, and displayed to assess filtered and selected crosslinks in the context of the physical protein structure. Finally, filtered sets of cross-links and individual views can be exported.

**Figure 1.**
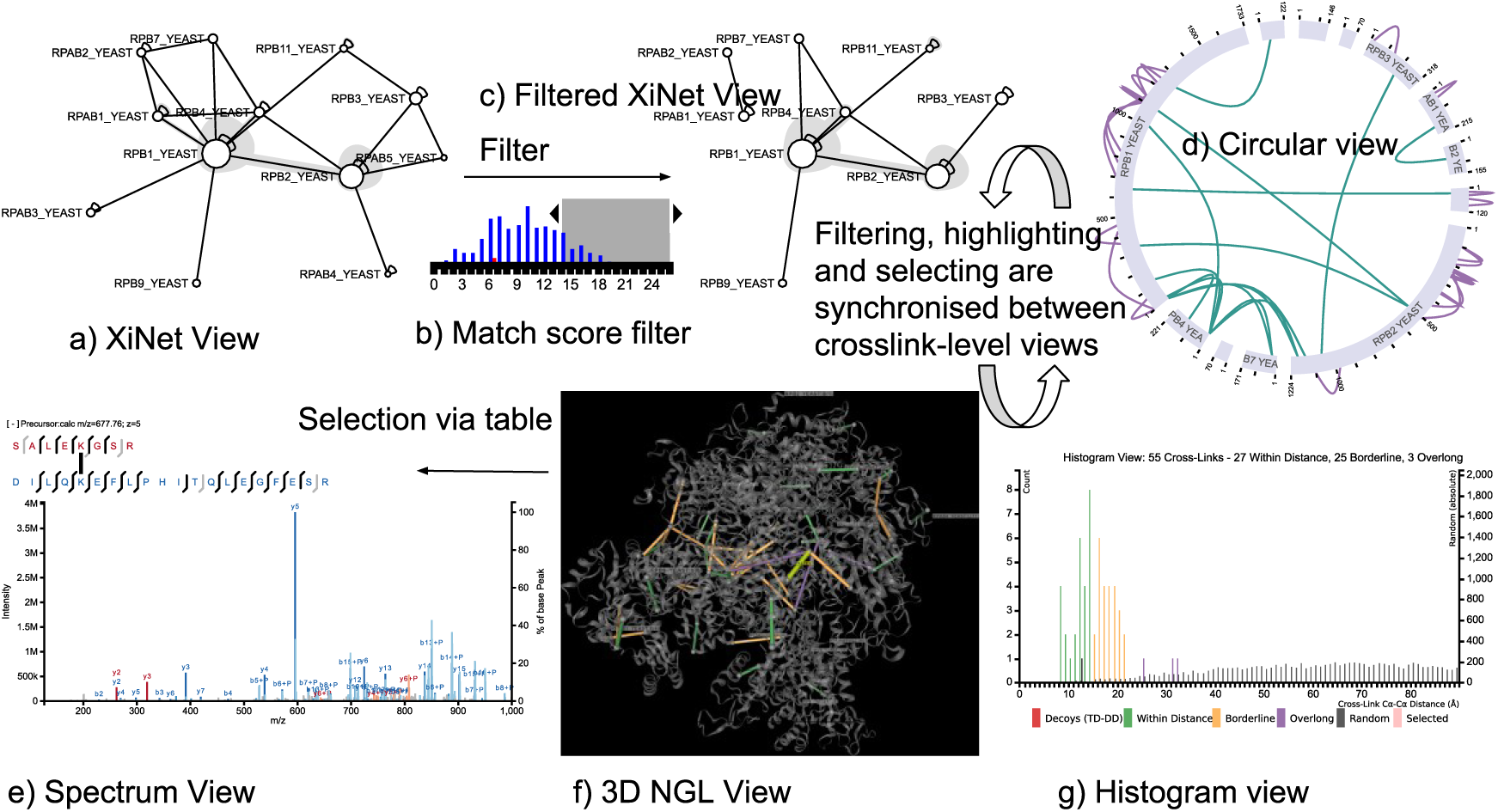
Some of the interactive components of XiView. Clockwise from top left: Initial network view, b) visual filter for match scores, c) the resulting filtered network view, d) circular view with same filtering. Upon loading of a PDB file we can see g) a histogram view showing distance against random background with a distance colour scheme, f) a 3D NGL view with crosslinks automatically aligned to PDB sequences and a distance colour scheme. Finally we can interrogate specific PSM details in e) the spectrum view. Selected links populate a table of PSM’s and spectra is launched from there. Views a) d) f) and g) all respond immediately to changes in the filter, along with user selections and highlighting.

xiView’s capabilities have evolved according to the demands of a set of real scientific users and have also undergone testing to discover and correct usability issues, and thus can be confidently said to support real user tasks in inspecting crosslinking data sets.

Documentation for the software can be found at https://xiview.org/xidocs/html/xiview.html, and short tutorial videos can be found at http://rappsilberlab.org/rappsilber-laboratory-home-page/tools/xiview/xiview-videos/.

A public facing version of the software is available at https://xiview.org where example data sets can be explored and new datasets can be uploaded and explored. There is a file size limit of 1GB for uploads. For datasets with a file sizes larger than 1GB, or where users do not wish their data stored by a third party, instructions for a local installation of xiView are available at https://github.com/xiView_container.

## Code availability

Source code is available at https://github.com/xiView_container. This code is published under the open source license GPL v3.

## Acknowledgments

We acknowledge the work of Alex Rose whose NGL 3D viewer^15^ is embedded as the basis of one of the views within xiView and Lutz Fischer, the author of the xiAnnotator which xiSPEC^13^ uses to annotate the spectra. Thanks to Zhuo Chen for the use of the example data set. This work was supported by the Wellcome Trust through a Senior Research Fellowship to J.R. [103139]. The Wellcome Centre for Cell Biology is supported by core funding from the Wellcome Trust [203149].

## Author Contributions

M.G., C.C., and L.K. wrote the software, J.R. conceived of the project. All authors contributed to writing the manuscript.

